# Chronic exposure of ErbB TKIs inhibits DRA expression and activity through an ERK/Elk-1/CREB/AP-1 dependent pathway

**DOI:** 10.1101/2020.10.16.342139

**Authors:** Hong Yang, Ru-Xian Lin, Rafiquel Sarker, Mark Donowitz

**Affiliations:** Department of Medicine, Division of Gastroenterology and Hepatology, Johns Hopkins University School of Medicine, Baltimore, Maryland; Department of Physiology, Johns Hopkins University School of Medicine, Baltimore, Maryland

**Keywords:** the human erythroblastic oncogene B receptor tyrosine kinase inhibitor (ErbB TKI), diarrhea, down regulated in adenoma (DRA), cystic fibrosis transmembrane conductance regulator (CFTR), extracellular-signal-regulated kinase (ERK), ETS like kinase 1 (Elk-1), cAMP response element-binding protein (CREB), activating protein 1 (AP-1)

## Abstract

Diarrhea is the major side effect of first- and second-generation ErbB tyrosine kinase inhibitors (TKI), the mechanism of which remains incompletely understood. The current studies were carried out over the time frame that ErbB TKIs usually initiate diarrhea. We report in Caco-2/bbe cells that exposure of ErbB TKIs, but not non-ErbB TKIs for six days at clinically-relevant concentrations significantly reduced the expression of DRA and inhibited apical Cl^-^/HCO_3_^-^exchange activity. The ErbB TKIs decreased DRA expression through an ERK/Elk-1/CREB/AP-1 dependent pathway. The blockade of ERK phosphorylation by ErbB TKIs decreased the phosphorylation of Elk-1 and the amount of total and p-CREB, and reduced the expression of C-Fos, which is part of the AP-1 complex that maintain DRA expression. Altogether, our studies demonstrate that ErbB TKIs decrease expression and activity of DRA, which occurs over the time frame that these drugs clinically cause diarrhea, and since DRA is part of the intestinal neutral NaCl absorptive process, the reduced absorption is likely to represent a major contributor to the ErbB TKI-associated diarrhea.

## Introduction

The human erythroblastic oncogene B (ErbB) receptor belongs to the receptor tyrosine kinase (RTK) family, which include four members, ErbB1 to ErbB 4. The small molecule inhibitors that target the ErbB family of receptor tyrosine kinases (ErbB TKIs) have gained extensive clinical application for the treatment of ErbB-overexpressing carcinomas, especially breast cancer and non-small cell lung cancer (NSCLC) (Roskoski, 2014). Diarrhea is a major side effect of first- and especially second-generation ErbB TKIs and occurs in 18% - 69% and 65% - 96% of treated patients, respectively, and can lead to dose reduction, interruption, or even discontinuation of treatment (Rugo et al, 2019). The currently available management of ErbB TKI-associated diarrhea is mainly symptom-oriented, including fluid and electrolyte replacement to treat dehydration, and loperamide (Hirsh et al, 2014), which is of limited effectiveness especially for treating severe diarrhea (Carlos Hernando Barcenas, 2019). Therefore, there remains an unmet need to understand the mechanism(s) and develop efficacious, targeted antidiarrheal therapy directed to ErbB TKIs.

Physiologically, the movement of Na^+^, K^+^, Cl^-^ and HCO_3_^-^ between the intestinal lumen and blood is regulated by specific transport proteins that are differentially distributed along the horizontal and vertical axes of the intestine. Most diarrheal diseases are caused by excessive secretion and/or reduced absorption (Field, 2003). Intestinal fluid secretion is driven by active chloride ion (Cl^-^) secretion from the blood to the intestinal lumen. The Na^+^-K^+^-2Cl^-^ symporter (NKCC) in the basolateral membrane transports Cl^-^ from the blood into the enterocytes via the Na^+^ concentration gradient created by the Na^+^/K^+^-ATPase. The apical Cl^-^ channels, mainly the Ca^2+^-activated Cl^-^ channels (CaCCs) and cyclic-nucleotide-activated cystic fibrosis transmembrane conductance regulator (CFTR) allow Cl^−^ secretion into the intestinal lumen by using the interior-negative cell potential maintained by basolateral K^+^ channels and the Cl-concentration gradient. Intestinal fluid absorption is driven primarily by active Na^+^ transport accompanied by Cl^-^ and HCO3^-^ absorption which occurs during the period between meals. It is an electroneutral process that involves the apical sodium/hydrogen exchanger 3 (NHE3), using the Na^+^ gradient created by the Na^+^/K^+^-ATPase, and the linked Cl /HCO_3_ exchanger, SLC26A3 (down regulated in adenoma (DRA)) and is probably contributed to by SLC26A6 (putative anion transporter-1 (PAT-1). In multiple diarrheal diseases, enterotoxins from bacteria, such as cholera toxin, E.coli heat labile enterotoxin and viruses such as rotavirus dramatically increase the intracellular levels of cAMP and/or Ca^2+^, respectively, which activate the apical Cl^-^ channels, CFTR and CaCC, respectively, and the basolateral K^+^ channels, and inhibit the Na^+^ absorptive proteins NHE3 and/or DRA, contributing to secretory diarrhea (Das et al, 2018; Hodges & Gill, 2010). In addition, NHE3 and DRA mutations with reduced or absent transporter activity are causally related to congenital human Na^+^- and Cl^-^-losing diarrheal disorders (Hoglund et al, 1996; Janecke et al, 2015). In addition, loss of DRA expression has been implicated as a crucial contributor to *Salmonella-induced* diarrhea (Barrett, 2020).

Although multiple mechanisms have been suggested as contributing to ErbB TKI-associated diarrhea, blocking of the inhibitory effect of the EGF signaling pathway on Cl^-^ secretion has been accepted as the primary mechanism (Barrett, 2020; Hirsh et al, 2014; Kim et al, 2020; Rugo et al, 2019; Uribe et al, 1996; Van Sebille et al, 2015). The recent publication by Duan et al indicated that ErbB TKIs cause diarrhea by activating basolateral K^+^ channels and apical CFTR, which would be otherwise be inhibited by EGF signaling through PKC and ERK dependent pathways (Duan et al, 2019). However, the mechanism of TKI diarrhea remains only partially understood, since these previous studies were performed in a way that only evaluated acute effects of interfering with EGF signaling on Cl^-^ secretion, while ErbB TKI-associated diarrhea usually occurs in a more delayed manner, with onset typically starting days or weeks after initiation of the treatment (Chan et al, 2016; Crown et al, 2008; Hopkins et al, 2018; Rugo et al, 2019). We therefore evaluated the effect of ErbB TKIs on intestinal ion transport over the period that ErbB TKIs usually initiate diarrhea, and report studies using the differentiated and polarized *in vitro* intestinal Caco-2/bbe cell model.

## Results

### Chronic exposure to ErbB TKIs increases CFTR and decreases DRA expression in Caco-2/bbe cells

Since EGFR has been shown to be involved in setting rates of intestinal Cl^-^ secretion and neutral NaCl absorption, we initially evaluated the effects of ErbB TKIs on CFTR and DRA. In order to mimic the time course over which ErbB TKIs usually lead to diarrhea, we initially investigated the effect on protein levels of DRA and CFTR of chronic exposure (6 days) to five first-, second- or third-generation ErbB TKIs at their clinically relevant plasma concentrations (afatinib, 0.2 μM, gefitinib, 0.5 μM, lapatinib, 2.5 μM, neratinib, 0.2 μM, and osimertinib, 0.5 μM; https://www.accessdata.fda.gov/). We also included three non-ErbB TKIs (dasatinib, 0.1 μM, suntinib, 0.25 μM and imatinib, 2.5 μM) for comparison. As shown in Figure 1A, treatment with ErbB TKIs significantly increased the protein level of CFTR, and decreased the protein expression of DRA. Intriguingly, the effects of ErbB TKIs on CFTR and DRA were not observed with exposure to non-ErbB TKIs, suggesting an involvement of ErbB signaling in the regulation of CFTR and DRA expression. There was no significant alteration in TEER of the Caco2/-bbe monolayers after 6 days of TKI treatment (Figure 1B).

**Figure 1.**
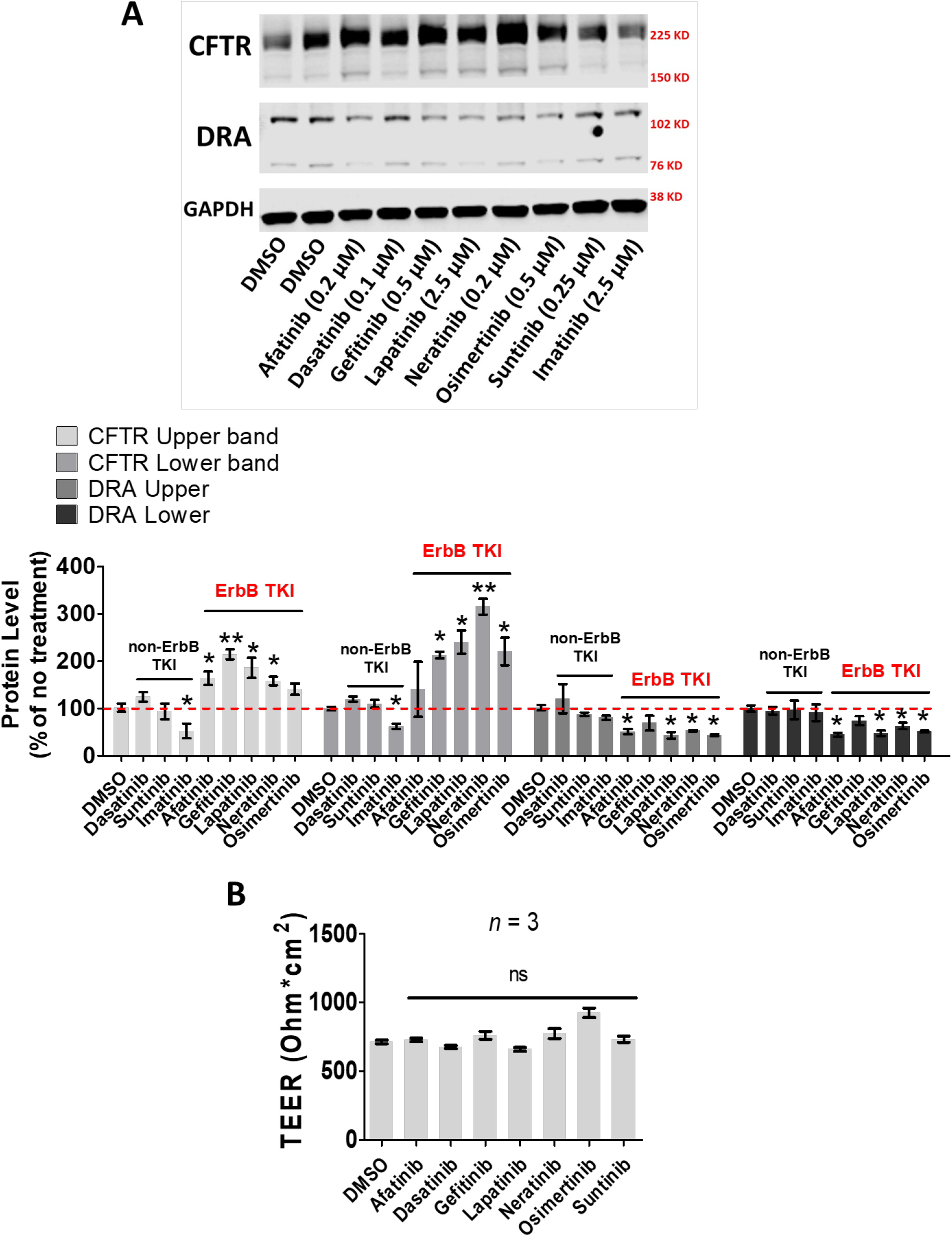
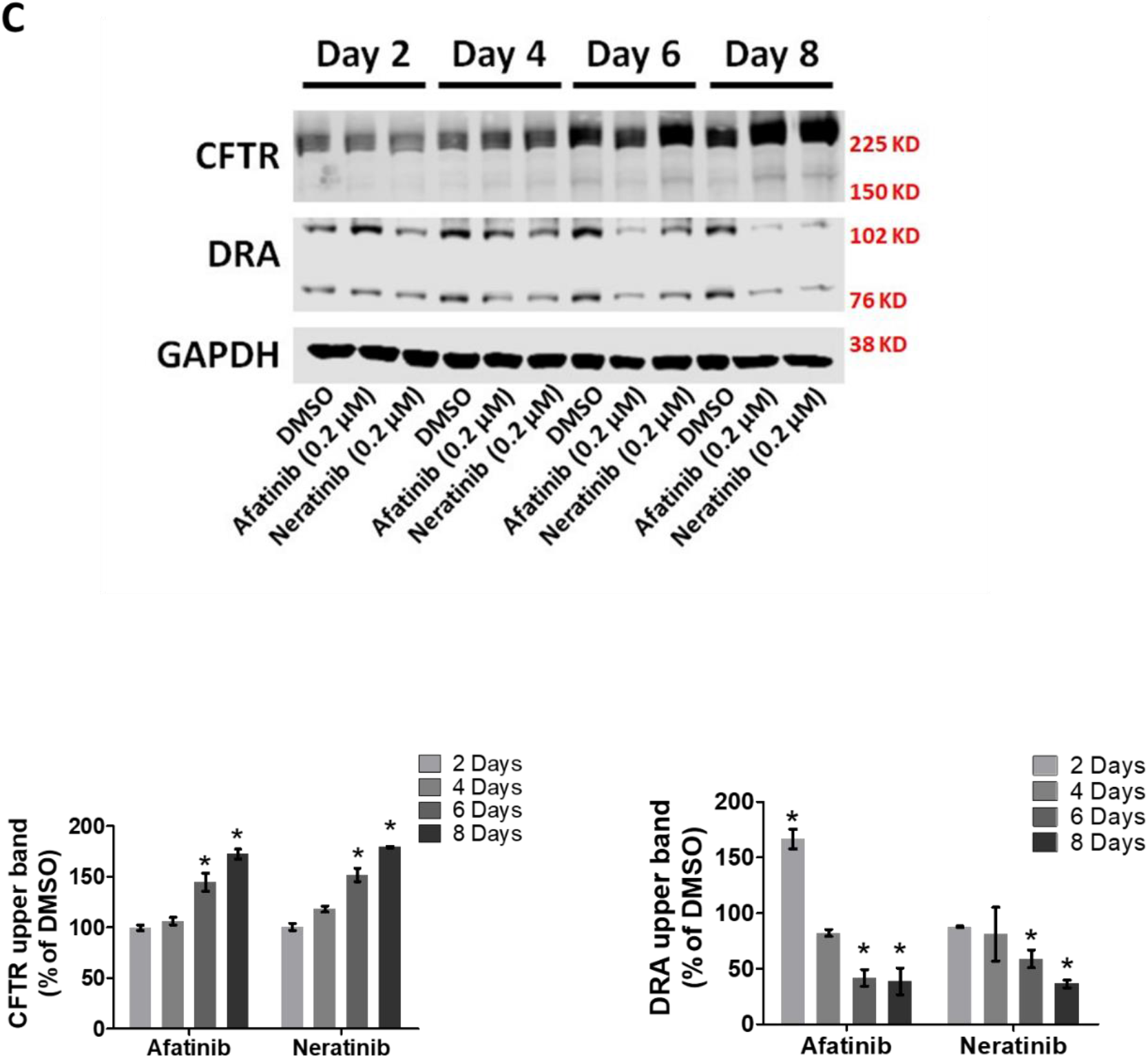
Chronic exposure of ErbB TKIs increases CFTR and decreases DRA expression in Caco-2/bbe cells. A, Representative immunoblotting of CFTR and DRA in Caco-2/bbe cells treated with several TKI drugs for 6 days, and quantification of the relative abundance of CFTR and DRA normalized to loading control in 3 independent experiments. B, Measurements of TEER in Caco-2/bbe cells treated with TKI drugs for 6 days. C, Representative immunoblotting of CFTR and DRA in Caco-2/bbe cells treated with indicated TKI drugs for 2 to 8 days, and quantification of the relative abundance of CFTR and DRA normalized to vehicle control, which is set as 100% in each of three independent experiments. Data presented as mean ± s.e.m. *, *p* < 0.05, **, *p* < 0.01, ns, not significant, as compared to untreated (A and B) or DMSO (C) (ANOVA followed by *post hoc* Tukey analysis).

We chose afatinib and neratinib to further investigate the time-dependent effects on CFTR and DRA expression in Caco-2/bbe cells (Figure 1C). Both afatinib and neratinib did not significantly alter CFTR and DRA expression during the first 4 days of treatment; however, 6 and 8 days of TKI exposure significantly decreased DRA expression. Likewise, six and 8 days of TKI exposure significantly increased CFTR expression. Since changes of CFTR and DRA protein expression were significantly altered by 6 days of treatment with ErbB TKIs, which aligns well with the time frame over which they initiate clinical diarrhea, we selected treatment for 6-days to be examined during the rest of the studies.

### ErbB TKIs alters the transcriptional expression of multiple ion transport proteins in Caco-2/bbe cells

Since chronic treatment with ErbB TKIs significantly affected the protein expression of both CFTR and DRA, we investigated effects on their mRNA expression as well as that of several additional transport proteins that are also involved in ion transport related to diarrhea (Figure 2). Consistent with the protein data, all five ErbB TKIs significantly increased the mRNA expression of CFTR and decreased that of DRA with a similar magnitude to the level of changes in protein expression. Intriguingly, treatment of ErbB TKIs also significantly increased the mRNA expression of basolateral K^+^ channels (KCNN2s, KCNN4 and KCNQ1) by ~3 fold, increased expression of NKCC1, although to a lesser extent, and decreased that of PAT-1 by ~70%. After we separately considered the data of all ErbB TKIs and separately that of all the non-ErbB TKIs, by taking the data for each TKI as *n* = 1, as shown in figure 2B, it was clear that the changes in transporter mRNA expression exclusively occurred with ErbB TKI exposure but not with the non-ErbB TKIs. Altogether, the effects of ErbB TKIs on DRA and CFTR protein and mRNA expression warranted further investigation of their functional alterations.

**Figure 2.**
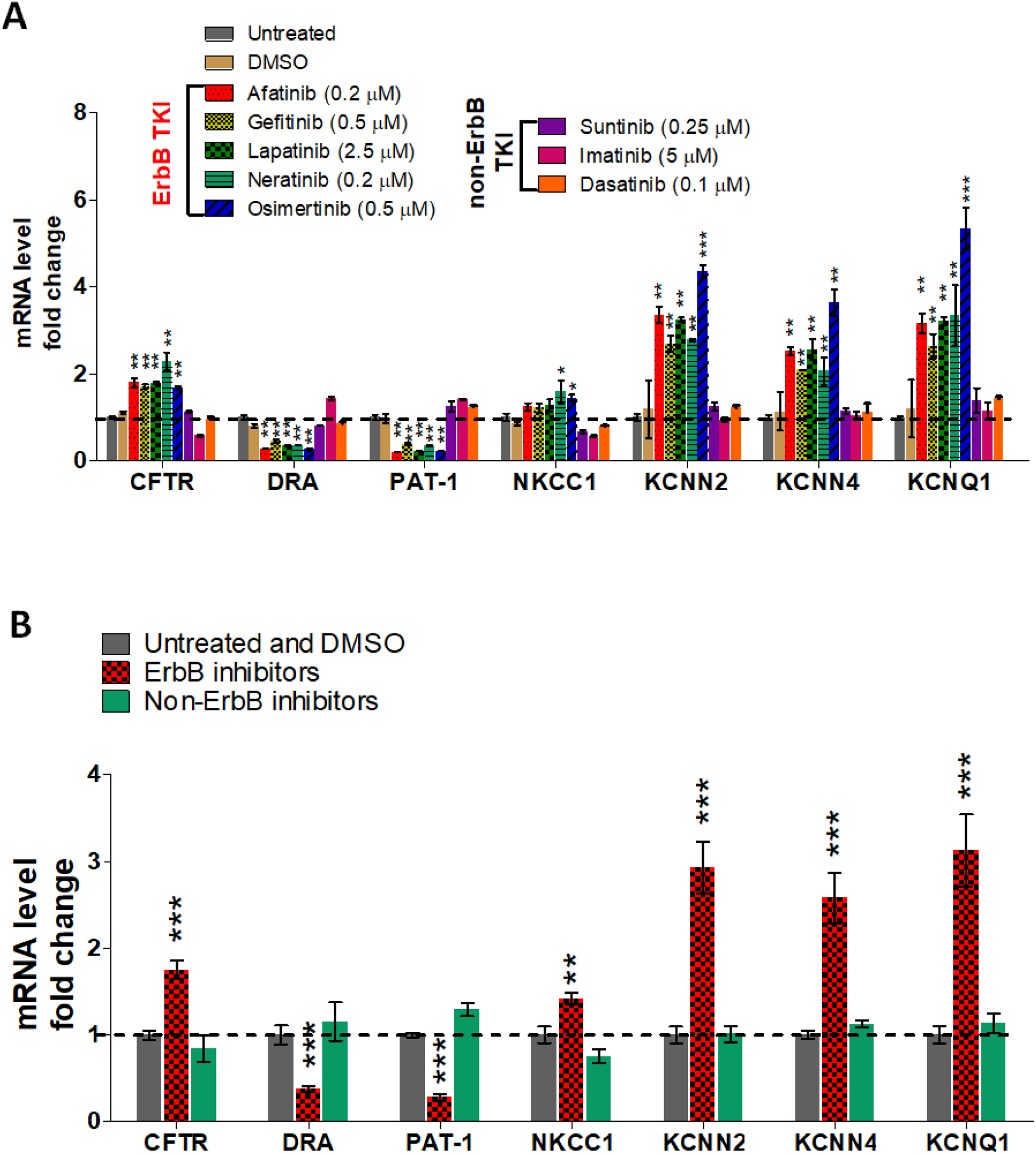
Chronic exposure of ErbB TKIs alters the transcriptional expression of transport proteins in Caco-2/bbe cells. A, mRNA expression of ion transport proteins in Caco-2/bbe cells treated with indicated TKI drugs for 6 days. B, mRNA expression of several ion transport proteins in Caco-2/bbe cells under three conditions (Untreated and DMSO, ErbB TKIs and Non-ErbB TKIs, *n* = 4, 5 and 3, respectively). Data presented as mean ± s.e.m. *, *p* < 0.05, **, *p* < 0.01, ***, *p* < 0.001, as compared to untreated (A) or untreated and DMSO (B) (ANOVA followed by *post hoc* Tukey analysis).

### ErbB TKIs diminish the forskolin-stimulated Isc in Caco-2/bbe cells

The upregulated expression of basolateral K^+^ channels, NKCC1 and CFTR suggested that TKIs would cause an increase in cAMP stimulated Cl^-^ and fluid secretion. Thus the forskolin (FSK)-stimulated shortcircuit current (Isc) was determined in Caco-2/bbe monolayers after 6-days of TKI exposure. The Isc response to FSK is recognized as primarily CFTR-dependent, which was further confirmed using the specific CFTR inhibitor, CFTR_inh_-172, which was added to the monolayers after FSK treatment. However, unexpectedly, chronic exposure of ErbB TKIs (afatinib and gefitinib) significantly lowered the I_sc_ response to FSK as compared to DMSO and dasatinib (Figure 3). A possible explanation is provided in the ‘Discussion’.

**Figure 3.**
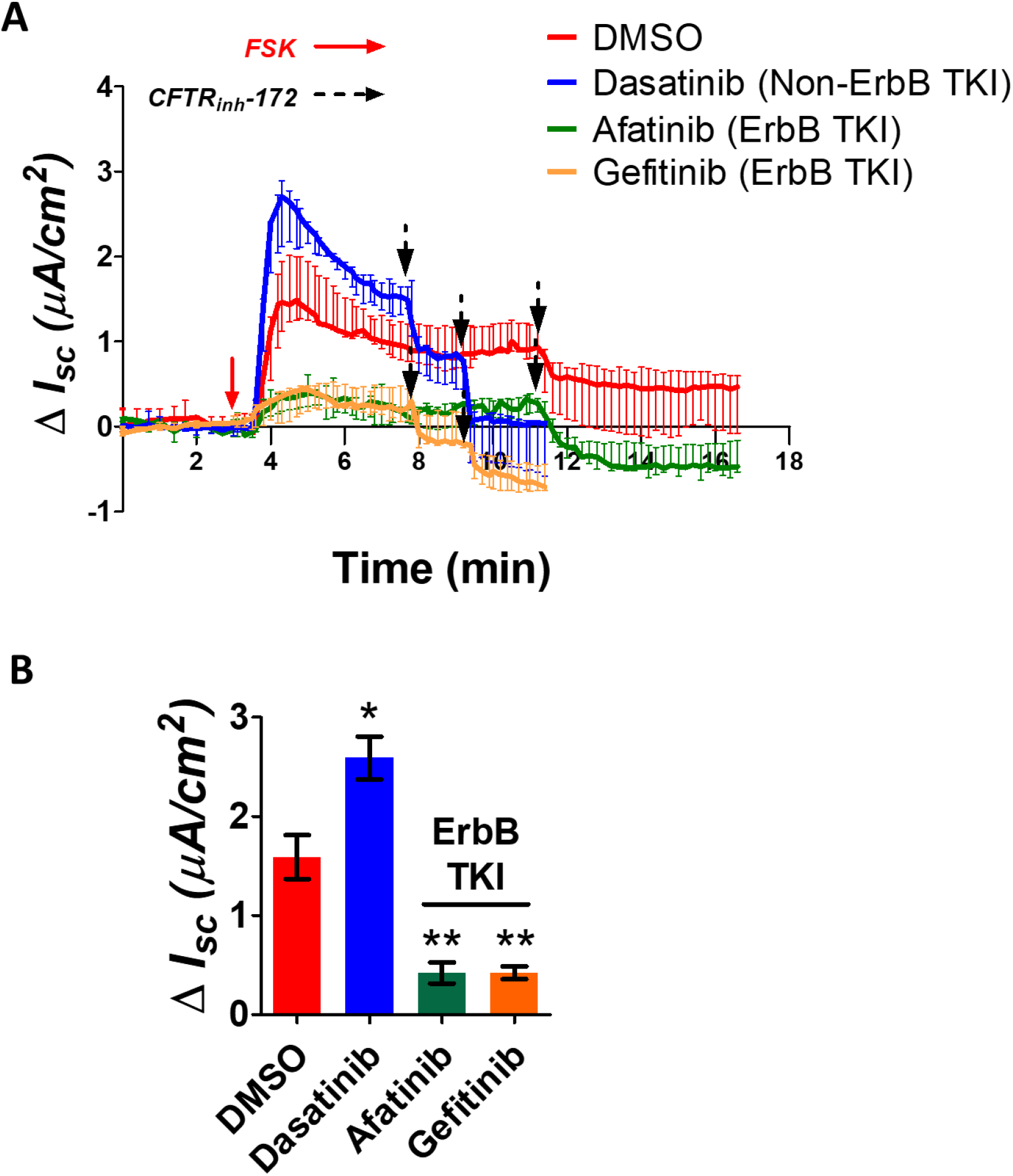
ErbB TKIs diminish the forskolin-stimulated short-circuit current in Caco-2/bbe cells. A, Change in Short-circuit current (Isc) from baseline in Caco-2/bbe cells showing response to Forskolin (FSK, 10 μM, red arrow) after treatment with indicated TKIs for 6 days. Black arrow indicates when CFTR_inh_-172 (10 μM) was added to the apical chamber. B, Quantification of peak I_sc_ response of figure 3A. Data presented as mean ± s.e.m. *n* = 3. *, *p* < 0.05, **, *p* < 0.01, as compared to DMSO (ANOVA followed by *post hoc* Tukey analysis).

### ErbB TKIs inhibit Cl^-^/HCO_3_^-^ exchange activity in Caco-2/bbe cells

The effect of chronic exposure of ErbB TKIs was investigated on Cl^-^/HCO_3_^-^ exchange activity, which is nearly entirely due to DRA activity in Caco-2/bbe cells. DRA activity was quantified as the extent of intracellular alkalinization initiated by extracellular Cl^-^ removal, and then exit of intracellular Cl^-^ in exchange for HCO_3_^-^ entry. This process is entirely inhibited in these cells by the specific DRA inhibitor, DRA_inh_-A250, as reported in our previous publication (Tse et al, 2019). As shown in figure 4A, a rapid intracellular alkalinization occurred immediately after removal of Cl^-^ from the apical surface of Caco-2/bbe cells in both vehicle (DMSO) and non-ErbB TKIs (dasatinib and suntinib) treated cells. In contrast, much slower alkalinization occurred with exposure to each of the ErbB TKIs (afatinib, gefitinib, lapatinib and neratinib). The initial rate of alkalinization (ΔpH_i_/min) was quantified in figure 4B to compare the effects on DRA activity. No significant differences were observed in non-ErbB TKIs compared to vehicle control. In contrast, much slower rates occurred in ErbB TKIs, particularly in those that belong to the second generation of TKIs (afatinib, reduce by 63.8 ± 7.0%; lapatinib by 70.2 ± 3.2% and neratinib by 66.1 ± 6.1%). Notably, the magnitude of the changes of activity aligns well with that of the decreases in protein and mRNA expression, as shown in Figures 1A (afatinib by 49.2 ± 7.8%; lapatinib by 57.0 ± 9.5% and neratinib by 48.7 ± 3.2%) and 2A (afatinib by 73.4 ± 2.1%; lapatinib by 66.2 ± 1.8% and neratinib by 64.6 ± 1.5%). Notably, gefitinib showed a trend to a smaller effect on DRA activity (by 43.5 ± 12.8%) as compared to the second-generation ErbB TKIs studied, even though statistical significance was not reached.

**Figure 4.**
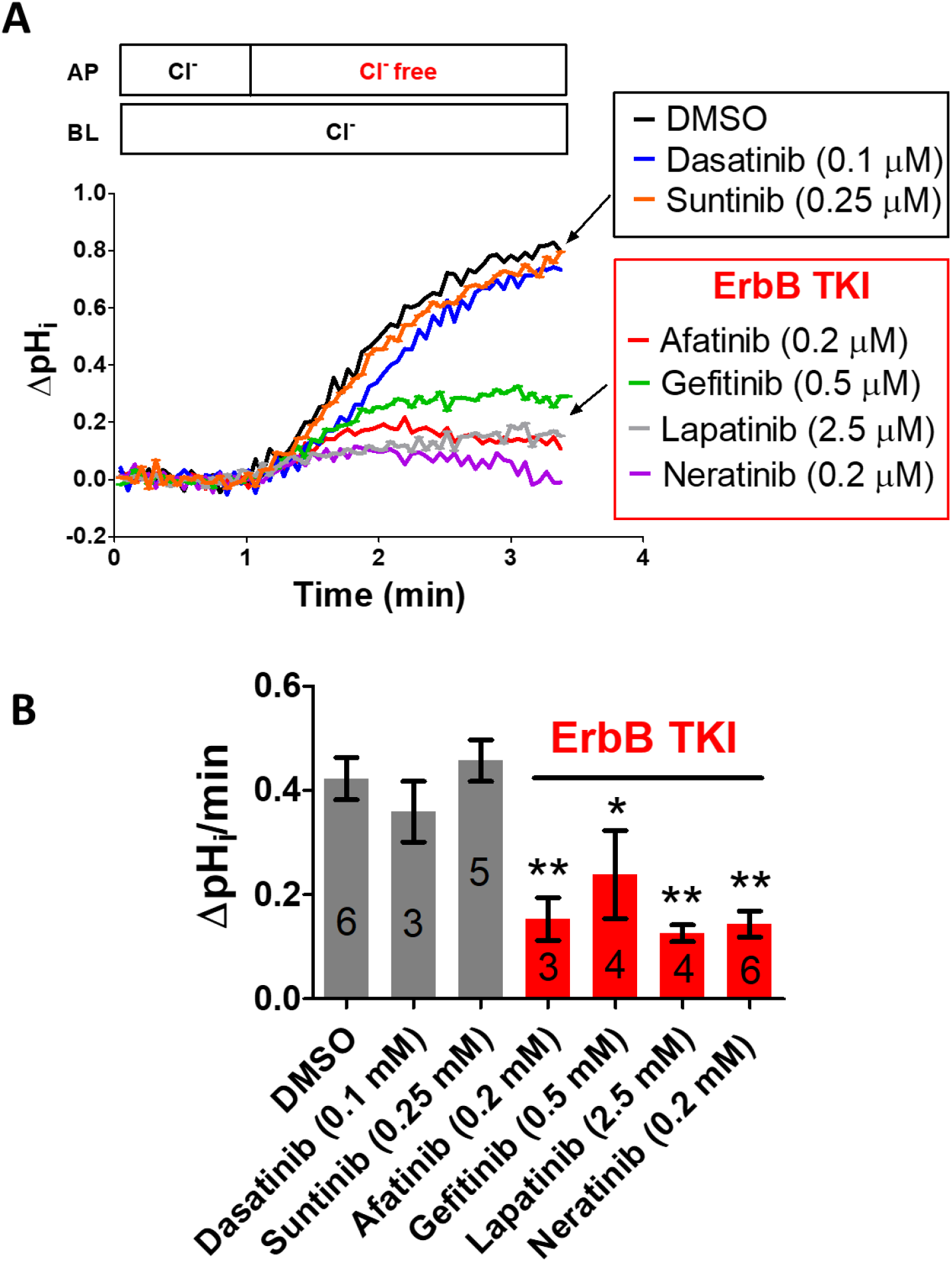
Chronic exposure of ErbB TKIs inhibits Cl^-^/HCO_3_^-^ exchange activity in Caco-2/bbe cells. A, Representative traces of Cl^-^/HCO_3_^-^ exchange are shown for Caco-2/bbe cells treated with indicated TKI drugs for 6 days. B, Initial Cl^-^/HCO3^-^ exchange rates are quantified for Caco-2/bbe cells treated with indicated TKI drugs in figure 4A. The number of experiments used in quantification is indicated inside each bar graph. Data presented as mean ± s.e.m. **, *p* < 0.01, as compared to DMSO (ANOVA followed by *post hoc* Tukey analysis).

### Extracellular-signal-regulated kinase (ERK) phosphorylation is involved in ErbB TKI-mediated loss of DRA expression and activity in Caco-2/bbe cells

We further investigated the cell signaling pathway(s) involved in ErbB TKIs-mediated loss of DRA expression. Signaling pathways including PLC-PKC, Ras-MAPK, PI3K-AKT are known to be prominent but not exclusive downstream effectors of ErbB signaling (Citri & Yarden, 2006; Hynes & MacDonald, 2009). Phosphorylation of PKC, ERK and AKT were selected as the main indicators of activity of each of these signaling pathways. Figure 5A shows that six days of exposure to ErbB TKIs significantly reduced the level of ERK phosphorylation (15.8% remaining compared to average of DMSO and non-ErbB TKIs), and to a much less extent, reduced phosphorylation of AKT (75.7% remaining) and PKC (81.9% remaining). To further clarify the role of each signaling pathway in regulation of DRA expression, specific inhibitors, including GDC-0994 (ERK inhibitor), RO.31.8220 (pan-PKC inhibitor) and MK-2206 (AKT inhibitor), were applied to Caco-2/bbe monolayer for 6 days in a concentration-dependent manner (Figure 5B). MK-2206 and RO.31.8220 dose-dependently reduced phosphorylation of AKT and PKC, respectively; however, RO.31.8220 did not alter DRA expression while MK-2206 only altered DRA at the highest concentration studied. At a lower concentration (0.1 μM), that reduced AKT phosphorylation to a similar extent as afatinib, MK-2206 did not alter DRA expression. GDC-0994 failed to inhibit ERK phosphorylation even at the highest concentration used (Germann et al, 2017), and also had no effect on DRA expression. In contrast, another ERK inhibitor (SCH772984), inhibits ERK phosphorylation, as reported (Morris et al, 2013) (Figure 5C). SCH772984 reduced DRA expression in a concentrationdependent manner, and at the highest concentration studied (5 μM) caused a comparable inhibition and reduction in expression of DRA as did afatinib (Fig 5D). These results support that ERK phosphorylation is a major regulator of DRA expression in Caco-2/bbe cells.

**Figure 5.**
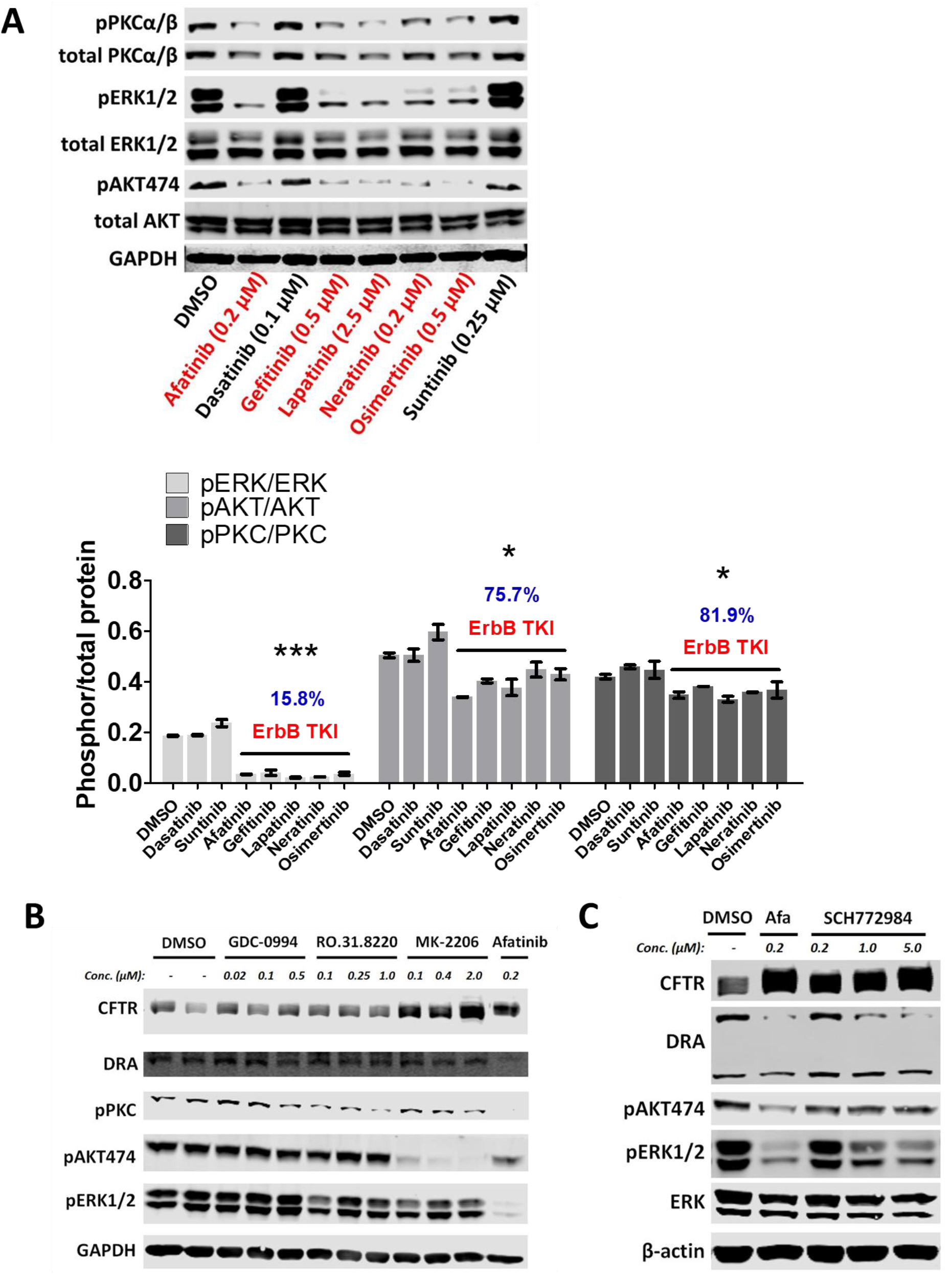
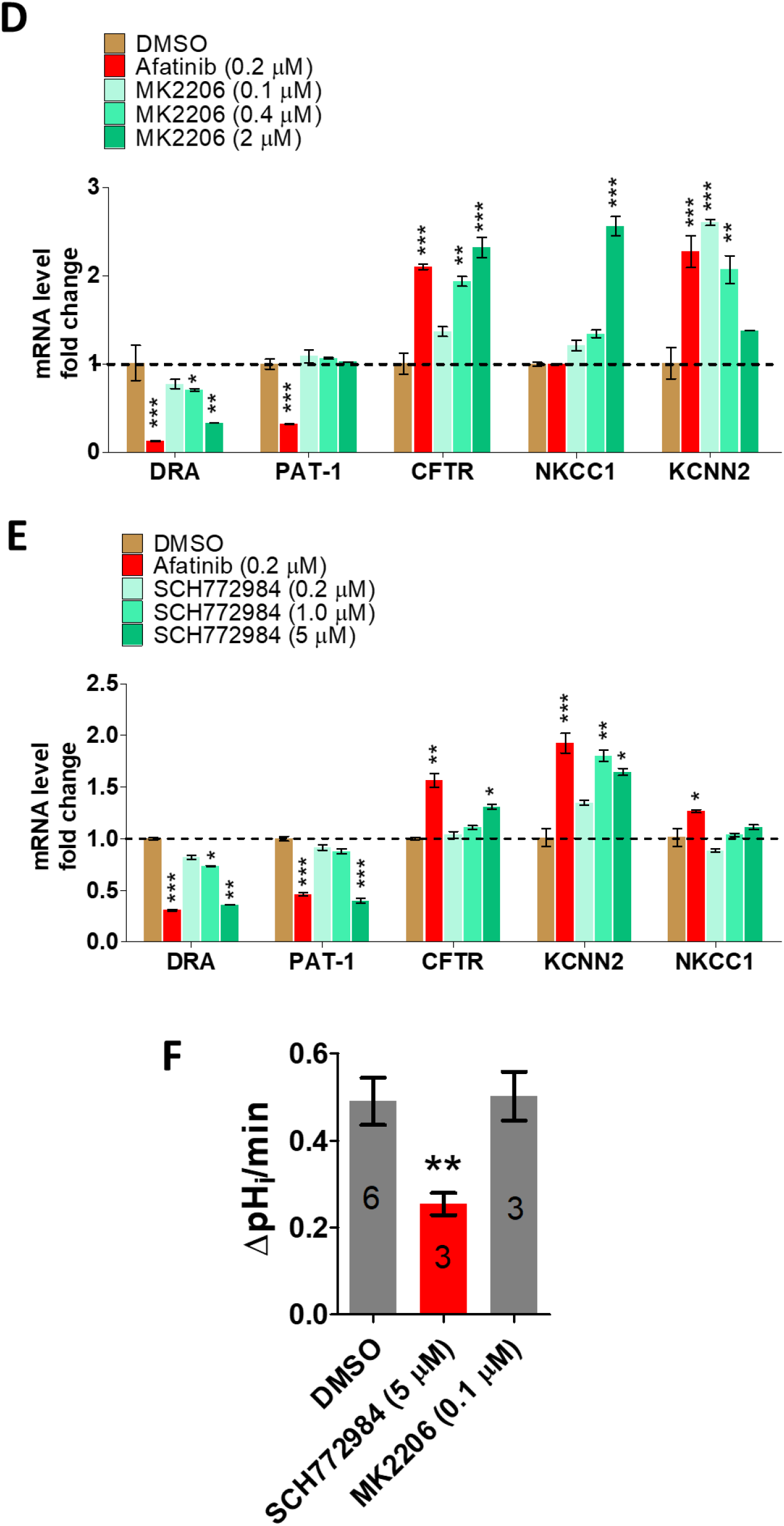
ERK phosphorylation is involved in ErbB TKIs-mediated loss of DRA expression in Caco-2/bbe cells. A, Representative immunoblotting of PKC, ERK and AKT in Caco-2/bbe cells treated with different TKI drugs for 6 days (left), and quantification of the relative abundance of phosphorylated protein normalized to total protein from 3 independent experiments (right). B, Representative immunoblotting of Caco-2/bbe cells treated with ERK inhibitor (GDC-0994), PKC inhibitor (RO.31.8220), AKT inhibitor (MK-2206) or afatinib at indicated concentrations for 6 days. Similar results were found in 3 independent experiments. C, Representative immunoblotting of Caco-2/bbe cells treated with ERK inhibitor (SCH772984) for 6 days. Similar results were found in 3 independent experiments. D, mRNA expression of several ion transport proteins in Caco-2/bbe cells treated with AKT inhibitor (MK-2206) for 6 days. *n* = 3. E, mRNA expression of ion transport proteins in Caco-2/bbe cells treated with ERK inhibitor (SCH772984) for 6 days. F, initial Cl^-^HCO^3-^ exchange rates are shown for Caco-2/bbe cells treated with indicated compounds for 6 days. The number of experiments used in quantification is indicated inside each bar. Data presented as mean ± s.e.m. *, *p* < 0.05, **, *p* < 0.01, **, *p* < 0.001, as compared to DMSO and non-ErbB TKI (A, Student’s t-test), as compared to DMSO (D, E, F, ANOVA followed by *post hoc* Tukey analysis).

Studied in parallel to DRA, the expression of CFTR was not affected by RO.31.8220 but was increased in a concentration-dependent manner by MK-2206 exposure. These results implicate involvement of AKT in the inhibitory regulation of CFTR expression, which has been previously reported (Reilly et al, 2017). In addition, SCH772984 similarly caused a concentration-dependent increase of CFTR expression, which also supported a role of the ERK pathway in the negative regulation of CFTR expression, as reported previously (Estelle Cormet-Boyaka et al, 2016).

The effect of MK-2206 and SCH772984 were examined on the transcriptional expression of several other ion transporters in the same Caco-2/bbe cells used for the DRA and CFTR studies (Figure 5D, 5E). MK-2206 at 0.1 μM, significantly increased KCNN2 and NKCC1 expression but only at the highest concentration studied. In contrast, SCH772984 (5 μM) significantly reduced PAT-1 expression and increased that of CFTR and KCNN2.

The Cl^-^/HCO3^-^ exchange activity was measured in Caco-2/bbe cells after chronic exposure to SCH772984 (5 μM) and MK-2206 (0.1 μM). As shown in figure 5F, SCH772984 but not MK-2206 caused a similar reduction of DRA activity as did the ErbB TKIs (Figure 4), further supporting a role of ERK signaling in DRA regulation.

### ETS like protein 1 (Elk-1), cAMP-response element binding protein (CREB), and activator protein 1 (AP-1) are involved in ErbB TKIs-mediated loss of DRA expression and activity in Caco-2/bbe cells

The increase in phosphorylation of ERK is known to stimulate the phosphorylation of the cAMP-response element binding protein (CREB) at a conserved serine site (Ser-133), which then binds to and promotes the expression of target genes (Sanna et al, 2018; Song et al, 2005). However, previous publications and our own sequence analysis failed to identify a potential CREB binding motif (CRE site) in the human DRA gene (*SLC26A3*), suggesting CREB is not a direct regulator of DRA expression (Zhang et al, 2005). Importantly, we identified that two components of the AP-1 complex, C-Fos and C-Jun, contained CRE sites, the expression of which could be increased by pCREB (Gustems et al, 2014). Notably, it has also been reported that the C-Fos gene contains a serum response element (SRE) site (Piechaczyk & Blanchard, 1994). Phosphorylated ERK is known to promote phosphorylation of Elk-1 (pElk-1), which forms a complex with its co-factor serum response factor (SRF) to promote C-Fos expression by binding to the SRE site. AP-1 is a transcriptional factor that regulates expression of a variety of genes by binding to a consensus sequence 5’-TGAC/GTCA-3’, which is also named, the 12-O-Tetradecanoylphorbol-12-Acetate (TPA) response element (TRE site). We analyzed the gene sequences of human DRA (*SLC26A3*) and PAT-1 (*SLC26A6*), and identified TRE sites in both genes, as shown in figure 6A. We further showed, in figure 6B, that the ErbB TKIs but not non-ErbB TKIs significantly decreased the level of pERK/ERK (14.5 ± 1.2% remaining compared to average of DMSO and non-ErbB TKIs), pElk/Elk (57.5 ± 6.2% remaining), CREB (69.0 ± 7.5% remaining), pCREB (64.4 ± 5.8% remaining), C-Fos (10.1 ± 0.8% remaining) and C-Jun (40.2 ± 2.1% remaining). Notably, the ratio of pCREB/CREB was not changed significantly, since both CREB and pCREB decreased by a similar extent. The involvement of Elk-1 and CREB further illustrated in figure 6C in which it was shown that the ERK inhibitor SCH772984 dose-dependently reduced the level of CREB, phosphorylation of CREB and Elk-1, and also reduced the expression of C-Jun and C-Fos, an effect aligning well with the DRA loss shown in figures 5C and 5E.

**Figure 6.**
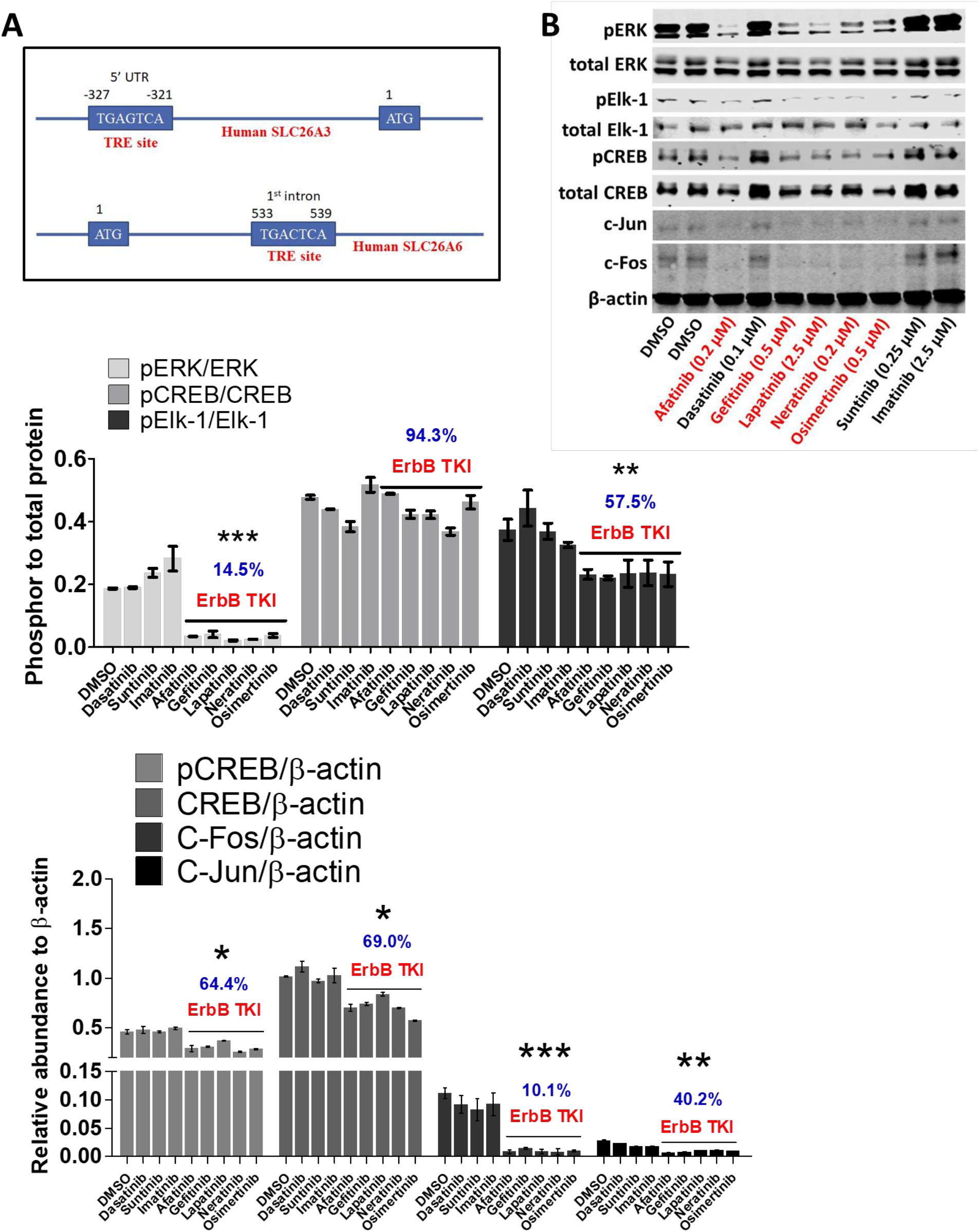
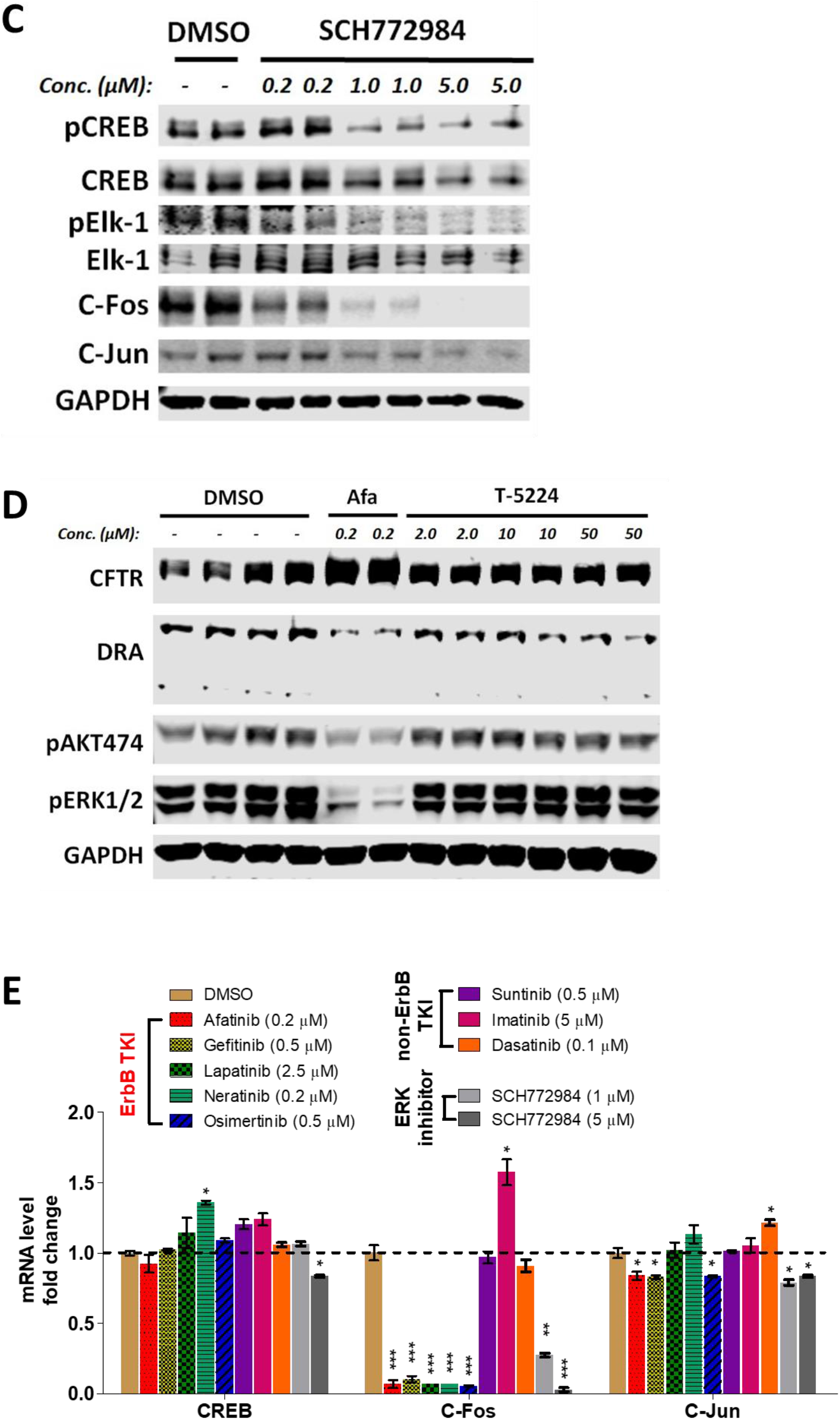
Elk-1, CREB and AP-1(C-Fos/C-Jun) are involved in ErbB TKIs-mediated loss of DRA expression in Caco-2/bbe cells. A, identification of TRE site for AP-1 (C-Fos/C-Jun) binding in human SLC26A3 and SLC26A6 genes. B, representative immunoblotting of pERK, total ERK, pCREB, total CREB, C-Jun and C-Fos in Caco-2/bbe cells treated with indicated TKIs for 6 days. Similar results were found in 3 independent experiments. Quantification was performed to calculate the ratio of phosphorylated to total protein (below left) and the relative abundance to β-actin (below right). C, Representative immunoblotting of Caco-2/bbe cells treated with ERK inhibitor (SCH772984) at indicated concentrations for 6 days. Similar results were found in 3 independent experiments .D, Representative immunoblotting of CFTR and DRA in Caco-2/bbe cells treated with AP-1 inhibitor T-5224 at indicated concentrations for 6 days. Similar results were found in 3 independent experiments. E, mRNA expression of CREB, C-Fos and C-Jun in Caco-2/bbe cells after treatment with indicated drugs for 6 days. *n* = 3. Data presented as mean ± s.e.m. *, *p* < 0.05, ***, p < 0.001, as compared to DMSO and non-ErbB TKI (A, Student’s t-test), as compared to DMSO (D, E) (ANOVA followed by post hoc Tukey analysis).

The involvement of AP-1 in DRA regulation was further indicated by use of the AP-1 specific inhibitor T-5224 (Yin, 2019). Figure 6D shows that T-5224 dose-dependently reduced the expression of DRA, but did not alter expression of pERK, pAKT and CFTR. The effect of TKIs and SCH772984 on transcriptional expression of CREB, C-Fos and C-Jun was also examined. As shown in Figure 6E, ErbB TKIs and SCH772984 dramatically reduced the expression of C-Fos, and reduced that of C-Jun to a less extent. Only SCH772984 at the highest concentration slightly but significantly reduced CREB mRNA.

## Discussion

The chronic manifestations of ErbB TKI-associated diarrhea and the crucial role of ion transporters played in the pathogenesis of diarrhea prompted us to investigate the effect of ErbB TKIs on epithelial ion transport over the time frame that ErbB TKIs usually initiate diarrhea. In this study, we report that chronic exposure of ErbB TKIs rather than non-ErbB TKIs for six days at clinically relevant concentrations significantly reduced the expression of DRA and its Cl7HCO_3_^-^ exchange activity in Caco-2/bbe monolayer *in vitro*. We showed that the mechanism by which ErbB TKIs decrease DRA expression is through an ERK/Elk-1/CREB-AP-1 (C-Fos/C-Jun) dependent pathway. The ErbB TKIs block ERK phosphorylation that decreases phosphorylation of Elk-1, reduces the amount of CREB and the corresponding amount of pCREB. This would be expected to reduce the amount of pElk-1 and pCREB binding to the SRE and CRE sites of AP-1 (C-Fos and C-Jun) respectively, and likely accounts for the reduced C-Fos and C-Jun demonstrated, which otherwise bind to a TRE site in the promoter of the DRA gene to maintain its expression. Our study, for the first time, investigated the chronic effects of ErbB TKIs on intestinal ion transporters *in vitro*, and identified loss of expression and function of DRA as a potential mechanism for ErbB TKIs-associated diarrhea.

The importance of DRA in the pathogenesis of diarrhea has been proven in both animal and human studies. DRA is most abundantly expressed in human colon, and to a less amount in ileum and duodenum (https://www.jrturnerlab.com/Transporter-Images), and is recognized as a major contributor to Cl^-^ and fluid absorption in the ileum and colon. Malfunctioning mutations in the human DRA (*SLC26A3*) gene have been etiologically linked to congenital chloride-losing diarrhea (CLD), an autosomal recessive disease featured by excessive chloride-loss in the stool, distention of the large intestine and life-long watery diarrhea (Hoglund et al, 1996; Holmberg et al, 1977; Kawamura & Nishiguchi, 2017). The clinical-pathological presentations of human CLD were successfully recapitulated in a Dra (*Slc26a3*) gene knockout mouse model, which exhibited similar diarrhea to that in patients with CLD, with decreased serum chloride level, volume depletion, and growth retardation (Schweinfest et al, 2006). While most of the *SLC26A3* mutations identified until now in CLD patients lead to almost non-functional DRA proteins (Makela et al, 2002), the ΔY526/7 mutation found in the STAS domain, does preserve ~50% of the activity (Dorwart et al, 2008). The mutation was first identified from a patient who was a heterozygote of the non-functional I675/6ins mutation and the ΔY526/7 mutation (Hoglund et al, 1998). Through a calculation of the gene-dose effect, a 75% loss of DRA activity appears severe enough to cause diarrhea. Notably, the second generation of ErbB TKIs, including afatinib, lapatinib and neratinib caused comparable DRA inhibition (close to 70%) in Caco-2/bbe cells (Figure 4), strongly implicating that DRA inhibition at least contributes to ErbB TKIs-associated diarrhea. In addition, the first-generation ErbB TKI gefitinib, which causes less severe diarrhea than second generation TKIs, exhibited a milder effect on DRA expression and function compared to the other second generation ErbB TKIs studied (Figure 1A, 2A and 4).

Our initial goal was to investigate the chronic effect(s) of ErbB TKIs on epithelial ion transport. Other than DRA, we have identified that the mRNA expression of several other ion transporters were affected by chronic treatment of ErbB TKIs in Caco-2/bbe cells. ErbB TKIs significantly increased both the transcriptional and translational expression of CFTR, and also the transcriptional expression of NKCC1 and basolateral K^+^ Channels (Figure 1A and 2A), which are necessary for intestinal anion secretion. However, contrary to our expectation, chronic exposure of ErbB TKIs (afatinib and gefitinib) lowered the FSK-stimulated I_sc_ as compared to DMSO and dasatinib (Figure 3A). FSK-stimulated I_sc_ has been shown to represent CFTR-dependent Cl^-^ secretion, and use of the CFTR specific inhibitor CFTR_inh_-172 further confirmed this conclusion in our study. This intriguing and contradictory phenomenon might be explained by the interrupted interaction between CFTR and DRA. CFTR and DRA proteins are co-expressed in the luminal membrane of a large population of ileal lower villus and upper crypt cells and in some surface epithelial cells and upper crypt cells of human proximal colon (Tse et al, 2019). It has been reported that the FSK-stimulated activity (open probability) of CFTR is dramatically increased by six fold when DRA is present, which uses its STAS domain to bind to the phosphorylated R domain of CFTR (Ko et al, 2004). The dramatic loss of DRA seen with ErbB TKI treatment removes the DRA-dependent increase in cAMP stimulation of CFTR activity. However, cAMP stimulation of Cl^-^ secretion via CFTR occurs in multiple types of cells that lack DRA and a full explanation is lacking, not only for the failure of chronic ErbB TKI treatment to lead to increased CFTR-dependent Cl^-^ secretion given the increase in CFTR expression, but also to the extent of secretion being reduced. Taken together, our data obtained from Caco-2/bbe cells do not support that increase of CFTR related Cl^-^ secretion contributes significantly to chronic ErbB TKI-associated diarrhea.

The fact that diarrhea is a more common side effect of first-, and especially second generation pan-ErbB TKIs as compared to non-ErbB TKIs indicates that there is an important role for the inhibition of ErbB signaling in the pathogenesis of diarrhea and conversely that ErbB signaling is important for normal intestinal fluid and electrolyte transport. After binding to a variety of endogenous ligands, the ErbB receptors initiate downstream signaling cascades through Ras-MAPK, PI3K-AKT, Src-FAK, and other cell signaling pathways to regulate cell proliferation, differentiation, apoptosis and migration (Citri & Yarden, 2006; Hynes & MacDonald, 2009). We have provided evidence that ERK phosphorylation is involved in DRA regulation, which is consistent with the studies reported by Kumar et al (Kumar et al, 2014). They reported that DRA expression in intestinal epithelial cells could be stimulated by the probiotic bifidobacterium which acted via pERK. This observation also highlights the potential of using bifidobacterium to correct ErbB TKI-associated diarrhea, which warrants further evaluation. Study from the same research group also indicated the involvement of the PI3K-AKT pathway in regulation of DRA but not PAT-1 expression (Singla et al, 2010). Similarly, our data in figure 5D showed that high concentrations of an AKT inhibitor MK-2206 significantly reduced DRA mRNA expression but not that of PAT-1. However, we suggest that AKT inhibition was not a crucial mechanism in ErbB TKI-mediated loss of DRA expression, since the same level of AKT inhibition achieved by MK-2206 (0.1 μM) as that by afatinib did not alter DRA mRNA and protein expression (Figure 5A and 5D), or DRA activity (Figure 5F).

Our studies indicate that ERK-dependent phosphorylation of Elk-1 and CREB, acting via pElk-1, and pCREB-dependent AP-1 (C-Jun/C-Fos) expression appear to be the downstream regulators of DRA expression. Direct regulation of CREB on DRA was ruled out since we and others failed to identify a potential CREB binding motif in the DRA gene (Zhang et al, 2005). We have provided evidence to support the role of AP-1 in DRA regulation, including loss of C-Jun/C-Fos expression by ErbB TKIs, identification of an AP-1 binding site in the DRA promoter, and loss of DRA expression by an AP-1 inhibitor. However, this mechanism should be further examined experimentally to show direct binding and activation of AP-1 at the DRA gene (chromatin immunoprecipitation (ChIP) assays and luciferase reporter assays). The phosphorylation of ERK promotes phosphorylation of Elk-1, which forms a complex with its co-factor SRF to promote C-Fos expression by binding to the SRE site of C-Fos gene(Piechaczyk & Blanchard, 1994). In addition, our data showed that the level of ERK phosphorylation is necessary to maintain the level of phosphorylated CREB (Figure 6B and C), although this appears to be via changing the entire amount of CREB. It has been reported that phosphorylated ERK further activates 90 kDa ribosomal S6 kinase (RSK), which then phosphorylates and activates CREB to initiate target gene expression (Xing et al, 1996). However, we observed a proportional reduction in both total and phosphorylated CREB by ErbB TKIs and the ERK inhibitor SCH772984, implicating that with chronic ErbB TKI treatment, the role of phosphorylated ERK involves maintaining CREB expression and phosphorylation, and this occurs at a post-translational level, given the unchanged CREB mRNA level, although the mechanism has not been determined. Notably, other publications did not observe a similar effect of ERK inhibitors on CREB expression (Fu et al, 2019; Li et al, 2018), a discrepancy which might be explained by the shorter period (maximum 24 hours) of exposure in those studies as compared to ours (6 days).

Our study warrants further validation in normal human enteroids. Studies using human colon cancer cell lines, such as Caco-2, do not always mimic what occurs in normal human intestine concerning regulation of transport proteins. In contrast, human enteroids derived from different parts of human donors’ intestine will allow us to perform a more comprehensive and physiological investigation on the effect of chronic ErbB TKIs on epithelial ion and fluid transport (Donowitz et al, 2020; Foulke-Abel et al, 2016).

In conclusion, the present study provides evidence to support the notion that the pathogenesis of ErbB TKI-associated diarrheas involve reduced DRA expression and function and the delay in effects on DRA expression probably contributes to the delayed time course of ErbB TKI-associated diarrhea.

## Experimental procedures

### Antibodies and reagents

Primary antibodies were from Cell Signaling: Phospho-Erk1/2-Thr202/Tyr204 (#8544, rabbit monoclonal, 1:1000), Erk1/2 (#9102, rabbit polyclonal, 1:1000), Phospho-CREB-Ser133 (#9198, rabbit monoclonal, 1:1000), CREB (#9104, mouse monoclonal, 1:1000), Phospho-Akt-Ser473 (#9271, rabbit polyclonal, 1:1000), pan-Akt (#4691, rabbit polyclonal, 1:2000), Phospho-PKCα/β-Thr638/641 (#9375, rabbit polyclonal, 1:1000), PKCα (#59754, rabbit monoclonal, 1:1000); Santa Cruz Biotech: SLC26A3 (sc-376187, mouse monoclonal, 1:100), C-Fos (sc-166940, mouse monoclonal, 1:500), C-Jun (sc-74543, mouse monoclonal, 1:500); Sigma: GAPDH (G8795, mouse monoclonal, 1:1000), β-actin (A2228, mouse monoclonal, 1:1000); the Cystic Fibrosis Foundation Therapeutics: CFTR (#217, mouse polyclonal, 1:2000). Reagents were from Cayman Chemical: nigericin (#11437), GDC-0994 (#21107), SCH772984 (#19166), RO.31.8220 (#13334); MedChemExpress: T-5224 (HY-12270); Biovision: MK-2206 (1888-500); APExBIO: neratinib (A8322), CFTR_mh_-172 (B1435); AK Scientific: lapatinib (A817); Sigma: afatinib (A170920), dasatinib (SML2589), gefitinib (SML1657), osimertinib (A1577172), Suntinib (PZ0012), imatinib (SML1027).

### Cell culture

Caco-2/bbe cells (originally from M. Mooseker and separately J. Turner) were cultured in Dulbecco’s modified Eagle medium (10-017-CV) supplemented with 10% fetal bovine serum, 100 U/mL penicillin, and 100 mg/mL streptomycin in a humidified atmosphere with 5% CO_2_ at 37 °C. For experiments, cells were plated on Transwell inserts (Corning, Inc, Corning, NY, 0.4 μm pore, polyester) and studied at 10 –16 days after reaching confluency. For a typical 6-days treatment of ErbB TKIs, the drugs were applied to the monolayer from both apical and basolateral sides from day 10 to 16 after reaching confluency.

### Immunoblotting analysis

Cells were rinsed 3 times with phosphate-buffered saline, then solubilized in lysis buffer (60 mM HEPES, 150 mM NaCl, 3 mM KCl, 5 mM EDTA trisodium, 3 mM ethylene glycol-bis(b-aminoethyl ether)-N,N,N0,N0-tetraacetic acid, 1 mM Na_3_PO_4_, and 1% Triton X-100, 0.1% SDS, pH 7.4) containing a protease inhibitor cocktail. To avoid membrane protein aggregation, protein samples were incubated for 30 min at room temperature in Laemmli sample buffer before electrophoresis. All proteins were then transferred to a nitrocellulose membrane (0.45 μm). Membranes were blocked in 5% BSA for detection of phospho-proteins, while 5% milk was used for the other studies. After incubation with primary antibodies overnight at 4 °C, membranes were incubated against secondary antibodies (IRDye^®^ from Licor Inc.) at room temperature for 1 hour, then visualized and quantitated using an Odyssey CLx system and Image Studio software (LI-COR Biosciences, Lincoln, NE).

### Reverse transcription and real-time PCR

We followed a reported procedure to measure the transcriptional expression of target genes (Yin et al, 2018). In brief, the PureLink RNA Mini Kit (Life Technologies) was used to extract total RNA from Caco-2/bbe monolayers. Then the complementary DNA was synthesized from the extracted RNA using SuperScript VILO Master Mix (Life Technologies). Quantitative real-time PCR (qRT-PCR) was performed on a QuantStudio 12K Flex real-time PCR system (Applied Biosystems, Foster City, CA) by using Power SYBR Green Master Mix (Life Technologies). Samples were run in triplicate, and the relative mRNA expression level of targeted genes was calculated from the 2-ΔΔCT method normalized with the vehicle control and housekeeping β-actin RNA.

### Measurement of Cl^-^/HCO_3_^-^ exchange activity

A detailed procedure for measuring CT/HCO_3_^-^ exchange activity in Caco-2/bbe monolayers has been described previously (Tse et al, 2019). In brief, the activity was fluorometrically measured by using the pHi-sensitive dye BCECF-AM in a customized chamber allowing simultaneous but separate apical and basolateral superfusion. The monolayers were rinsed and equilibrated in sodium solution (138 mM NaCl, 5 mM KCl, 2 mM CaCl_2_, 1 mM MgSO_4_, 1 mM NaH_2_PO_4_, 10 mM glucose, 20 mM HEPES, pH 7.4) for 60 minutes at 37 °C, then loaded with BCECF-AM (10 mM) in the same solution for another 30 min, and mounted in a fluorometer (Photon Technology International, Birmingham, NJ). The basolateral surface was superfused continuously with Cl^-^ solution (110 mM NaCl, 5 mM KCl, 1 mM CaCl_2_, 1 mM MgSO_4_, 10 mM glucose, 25 mM NaHCO_3_, 1 mM amiloride, 5 mM HEPES, 95% O_2_/5% CO_2_), while the apical side was superfused with a shift betwen Cl^-^ solution or Cl^-^-free solution (110 mM Na-gluconate, 5 mM K-gluconate,5 mM Ca-gluconate, 1 mM Mg-gluconate, 10 mM glucose, 25 mM NaHCO_3_, 1 mM amiloride, 5 mM HEPES, 95% O_2_/5% CO_2_). The switch between Cl^-^ solution and Cl^-^-free solution results in HCO3^-^ uptake across the apical membrane is mediated by Cl^-^ /HCO_3_^-^ exchangers (such as DRA). The subsequent change of intracellular pH was recorded and the rate of initial alkalization was calculated using Origin 8.0 (Origin-Lab, Northampton, MA) as an indicator of Cl^-^/HCO_3_^-^ exchange activity.

### Measurement of short-circuit current

The short-circuit current was determined by the Ussing chamber/Voltage Clamp Technique (Tse et al, 2018). In brief, Caco-2/bbe monolayers were mounted in Ussing chambers and incubated in Krebs-Ringer bicarbonate (KBR) buffer (115 mM NaCl, 25 mM NaHCO_3_, 0.4 mM KH_2_PO_4_, 2.4 mM K_2_HPO_4_, 1.2 mM CaCl_2_, 1.2 mM MgCl_2_, pH 7.4) continuously gassed with 95% O_2_/5% CO_2_ at 37°C and connected to a voltage-current clamp apparatus (Physiological Instruments) via Ag/AgCl electrodes and 3 M KCl agar bridges. 10 mM glucose was supplemented as an energy substrate in the basolateral side, while 10 mM mannitol was added to the apical chamber to maintain the osmotic balance. Current clamping was employed and short-circuit current was recorded every 1 or 5 seconds by the Acquire & Analyze software 2.2.2 (Physiological Instruments, San Diego, CA, USA). 10 μM forskolin was added to the apical chamber to investigate cAMP-stimulated anion secretion. The specific CFTR inhibitor, CFTR_inh_-172 (10 μM), was added after forskolin to confirm the involvement of CFTR.

### Statistics

GraphPad Prism (version 6.01, GraphPad Software, San Diego, CA) was used to perform the statistical analysis. Data are mean ± s.e.m. of at least three independent experiments, with error bar equaling to one s.e.m. A two-tail Student’s t-test was used for statistical comparison between two groups, while one-way analysis of variance (ANOVA) followed by *post hoc Turkey* was adopted if more than two different groups were compared. *P < 0.05* was considered as the threshold of statistical significance.

**Figure 7.**
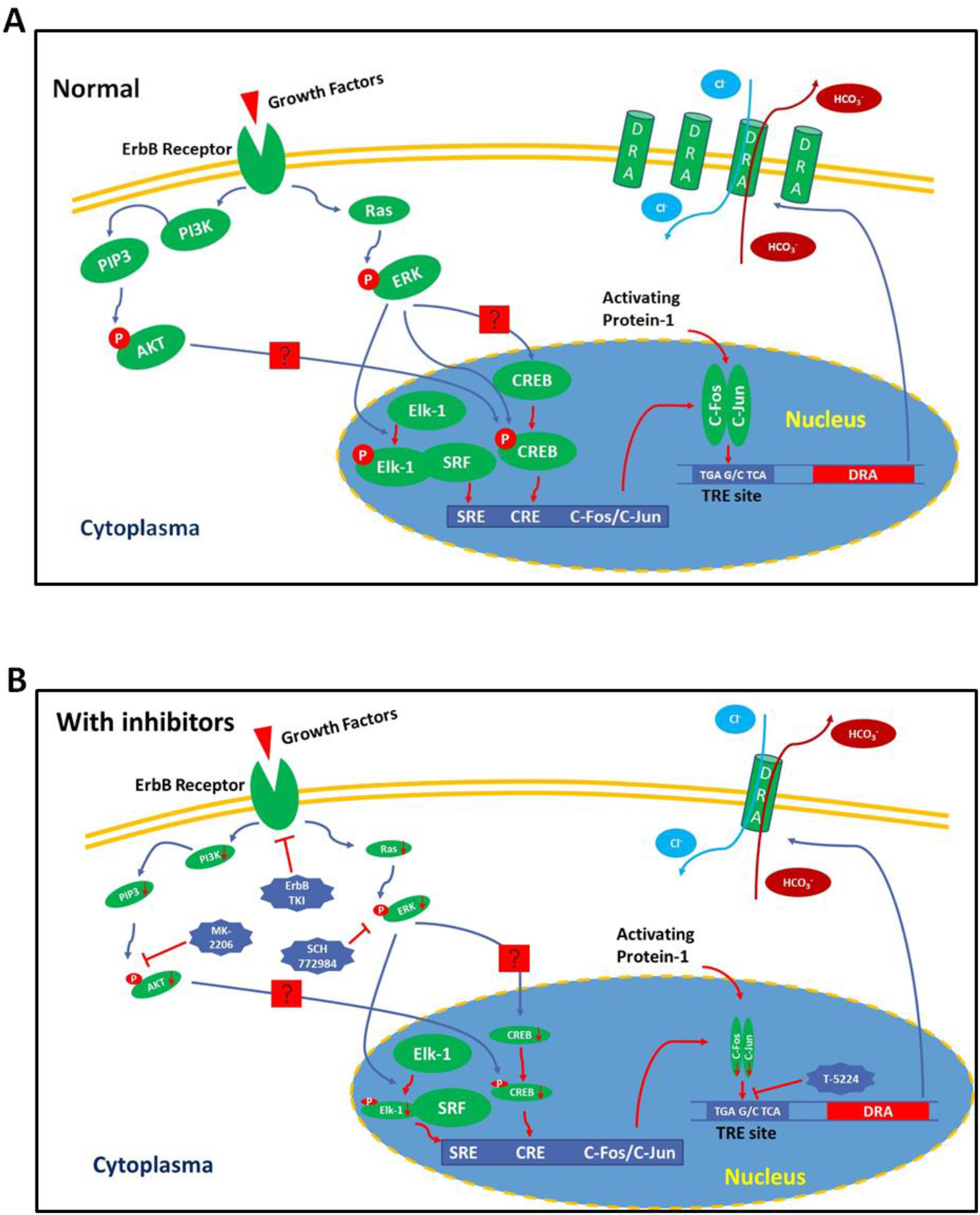
Schematic of mechanism of ErbB TKI on DRA expression. A, In normal state, growth factors bind to ErbB receptors on the cell membrane to initiate the signal cascades including Ras-ERK, PI3K-AKT. The Ras-ERK signaling pathways maintain CREB expression and stimulate phosphorylation of CREB (pCREB) and Elk-1 (pElk-1), then pElk-1/SRF and pCREB stimulate expression of C-Fos and C-Jun to form the activating protein-1 complex (AP-1), which binds to a TRE site in the promoter of DRA gene to maintain its expression. B, ErbB TKIs, ERK inhibitor SCH772984, AKT inhibitor MK-2206 and AP-1 inhibitor T-5224 block the signaling cascade to decrease DRA expression. SRE, serum response element; CRE, cAMP-response element; SRF, serum response factor; TRE, 12-O-Tetradecanoylphorbol-12-Acetate response elements.

## Data availability

All data are contained within the manuscript.

## Acknowledgments

We thank Dr. Alan S. Verkman (UCSF) and Dr. Shmuel Muallem (NIH), Dr. Chun-Ming Tse, Dr. Varsha Singh and Kelli Johnson for their helpful discussions and suggestions for this study.

## Author Contributions

HY and MD conceived and designed research. HY performed experiments, calculated results, interpreted data, prepared figures and drafted initial version of the manuscript. RS helped to modify transport assays used and helped to calculate and interpret results. RXL helped to develop and interpret the biochemical assays. HY and MD edited and revised manuscript. All authors read and approved the final version of manuscript.

